# Selective Postnatal Excitation of Neocortical Pyramidal Neurons Results in Distinctive Behavioral and Circuit Deficits in Adulthood

**DOI:** 10.1101/2020.01.18.911347

**Authors:** William E. Medendorp, Akash Pal, Madison Waddell, Andreas Björefeldt, Christopher I. Moore, Ute Hochgeschwender

**Author notes:** Correspondence (C.I.M.), (U.H.).

## Abstract

In leading models of Autism Spectrum Disorder, and in human data, the efficacy of outgoing cortical connectivity transitions from overly exuberant to languid from early development to adulthood. This transition begs the question of whether the early enhancement in excitation might be a common driver, across etiologies, of these symptoms. We directly tested this concept by chemogenetically driving neuronal activity in neocortical neurons during postnatal days 4-14. Hyperexcitation of Emx1-, but not dopamine transporter-, parvalbumin-, or Dlx5/6-expressing neurons led to decreased social interaction and increased grooming activity in adult animals. *In vivo* optogenetic interrogation in adults revealed decreased baseline but increased stimulus-evoked firing rates of pyramidal neurons, impaired recruitment of inhibitory neurons and reduced cortico-striatal communication. These results directly support the prediction that changed firing in developing circuits irreversibly alters adult circuit function that leads to maladaptive changes in behaviors. This experimental approach offers a valuable platform to study the impact of disruption of developmental neural activity on the formation and function of adult neural circuits and behavior.

## Introduction

While a variety of behavioral differences are associated with the heterogeneous phenotypes of adult Autism Spectrum Disorder (ASD), key cardinal features define this condition, including challenges with social skills, repetitive behaviors, speech and nonverbal communication (Lord *et al.*, 2000). Despite the existence of a final common ASD phenotype, it can be caused by a large variety of etiologies, from single gene mutations to metabolic, toxic, and social environmental insults (Watts, 2008; Chaste and Leboyer, 2012). ASD has been associated with over 250 genes, but with no single gene accounting for more than 1-2% of cases (Geschwind, 2011; Ramaswami and Geschwind, 2018). Commonality in adult expression of ASD suggests that a common mechanism(s) may be engaged through different etiologies.

Altered functional signaling through cortico-striatal projections (CS) is seen as a key potential driver of ASD symptoms (Fuccillo, 2016). Decreased CS connectivity is present in adult ASD patients (Abbott *et al.*, 2018; Balsters, Mantini and Wenderoth, 2018) and in several ASD mouse models (Peca *et al.*, 2011; Rothwell *et al.*, 2014; Wang *et al.*, 2016; Martella *et al.*, 2018; Shofty *et al.*, 2019). These specific changes are consistent with a broader reduction in MRI-based indicators of adult functional connectivity. However, Uddin et al. (Uddin, Supekar and Menon, 2013) found that while intrinsic functional connectivity in adults with ASD is generally reduced, functional connectivity in children with the disorder is increased. Interestingly, CS connectivity in postnatal development in a mouse model of autism (*Shank3B*−*/*−) also found increased CS network activity in development (Peixoto *et al.*, 2016, 2019) in contrast to the reduced CS functional connectivity in adult *Shank3* ASD models (Peca *et al.*, 2011; Wang *et al.*, 2016).

A key unanswered question posed by these studies is whether the transformation to weaker CS connectivity in the adult is related to the prior, stronger connectivity between projection pyramidal neurons to the striatum in development, or whether it is an independent process. In support of the former prediction, non-genetic insults during development that increase neocortical excitability, such as exposure *in utero* to the antiepileptic drug valproic acid, induce ASD phenotypes in both humans and mice (Roullet, Lai and Foster, 2013; Nicolini and Fahnestock, 2018), and in mice result in aberrant development of CS pathways (Kuo and Liu, 2017).

Further, a wealth of studies show altered excitatory-inhibitory (E/I) balance in the adult neocortex of ASD models, with a net shift to greater excitatory impact. Recent work analyzing multiple mouse models of ASD found that this net decrease in inhibitory efficacy can be relative, as excitatory and inhibitory synapses are both weakened (Antoine *et al.*, 2019). While acute changes in adult E/I balance can induce and reduce ASD phenotypes (Yizhar *et al.*, 2011; Selimbeyoglu *et al.*, 2017), whether these changes can also result from enhanced early life excitability in pyramidal neurons, or whether this shift has different etiological origin, is also unanswered.

To directly address the question of whether early life alterations in pyramidal activity can induce adult phenotypes and related circuit changes, we systematically enhanced pan-neocortical pyramidal activity levels in early development in healthy mice using BioLuminescent-OptoGenetic (BL-OG)-mediated chemogenetics (Berglund *et al.*, 2013, 2016; Moore and Berglund, 2019) to activate Emx1-positive neurons. We contrasted this activation with parallel activation of all neurons expressing the dopamine transporter (DAT), parvalbumin (Pvalb), or Dlx5/6. We found that developmental neocortical pyramidal over-drive in otherwise healthy subjects selectively led to decreased social interaction and increased grooming activity in adult animals, two key symptoms of ASD. Distinct, and in some cases opposed, behavioral deviations from control were observed in the other models. To test functional CS connectivity, we measured the impact of both endogenous spiking bursts and brief periods of optogenetically-induced pyramidal drive in the adult. Paralleling findings in adult Shank3 ASD mouse models, both drivers evoked significantly less cortically driven striatal activity, and these effects were paralleled by decreased cortico-striatal coherence in local-field potentials. Within neocortex, we also found decreased resting baseline activity but enhanced recruitment by optogenetic drive. This latter finding is well-explained by decreased feedback inhibition, suggesting diminished excitatory pyramidal output to local circuits in addition to the decrease in CS efficacy, as interneurons were more poorly recruited by this optogenetic pyramidal drive. These data show that brief and selective increases in pyramidal activity over a narrow developmental window can create both the behavioral and key neurophysiological signatures of ASD. As such, they suggest that a common driver of diverse insults is aberrant neocortical pyramidal activity early in life.

## Results

### Bioluminescent optogenetics allows developmental chemogenetic drive of distinct neural populations for adult optogenetic circuit interrogation and behavioral testing

To accommodate the need for non-invasive stimulation of genetically targeted neural circuits in a postnatally growing brain and temporally precise interrogation of the same circuits in the same animal as an adult, we employed bioluminescent optogenetics (BL-OG) (Berglund *et al.*, 2013, 2016; Park *et al.*, 2017; Song *et al.*, 2018; Zenchak *et al.*, 2018; Gomez-Ramirez *et al.*, 2019; Yu *et al.*, 2019). Here an optogenetic element can be activated either by light emitted from a tethered luciferase oxidizing a small molecule substrate, or by light emitted from physical sources, such as LEDs or lasers (Figure 1A). As we wanted to activate brain-wide genetically defined populations and do so starting soon after birth, we generated a conditional mouse line expressing the excitatory luminopsin 3 (LMO3) in a Cre dependent manner (Figures 1B and S1). Crossing this lox-stop-lox (LSL)-LMO3 line to various Cre driver mice enabled hyperexcitation of defined neuronal populations. We employed the Emx1-Cre line to enhance pan-neocortical pyramidal activity (Gorski *et al.*, 2002). Pvalb-Cre provided targeting to inhibitory neurons of the cortex (Hippenmeyer *et al.*, 2005), while Dlx5/6-Cre targets GABAergic neurons of the forebrain (Monory *et al.*, 2006). Lastly, DAT-Cre targets the dopaminergic system (Backman *et al.*, 2006), providing a line not expected to exhibit typical aspects of neurodevelopmental disorders such as social, anxiety, or depressive behaviors. Developmental hyperexcitation of these four different populations allowed us to test the impact of pyramidal cell hyperactivity while at the same time assessing whether early postnatal interference with neural activity would result in behavioral symptoms non-specifically, regardless of the population being activated, or whether activation of specific populations resulted in distinct behaviors that fall into phenotypic clusters (ASD, ADHD, anxiety, depression, etc.). By crossing heterozygous LSL-LMO3 mice to heterozygous Cre driver mice we generated extensive controls (Figure 1B). We used negative pups to control for genetic lineages and non-expressing littermates to control for any non-specific effects of administering the enabler of BL-OG, the luciferase substrate coelenterazine (CTZ).

**Figure 1.**
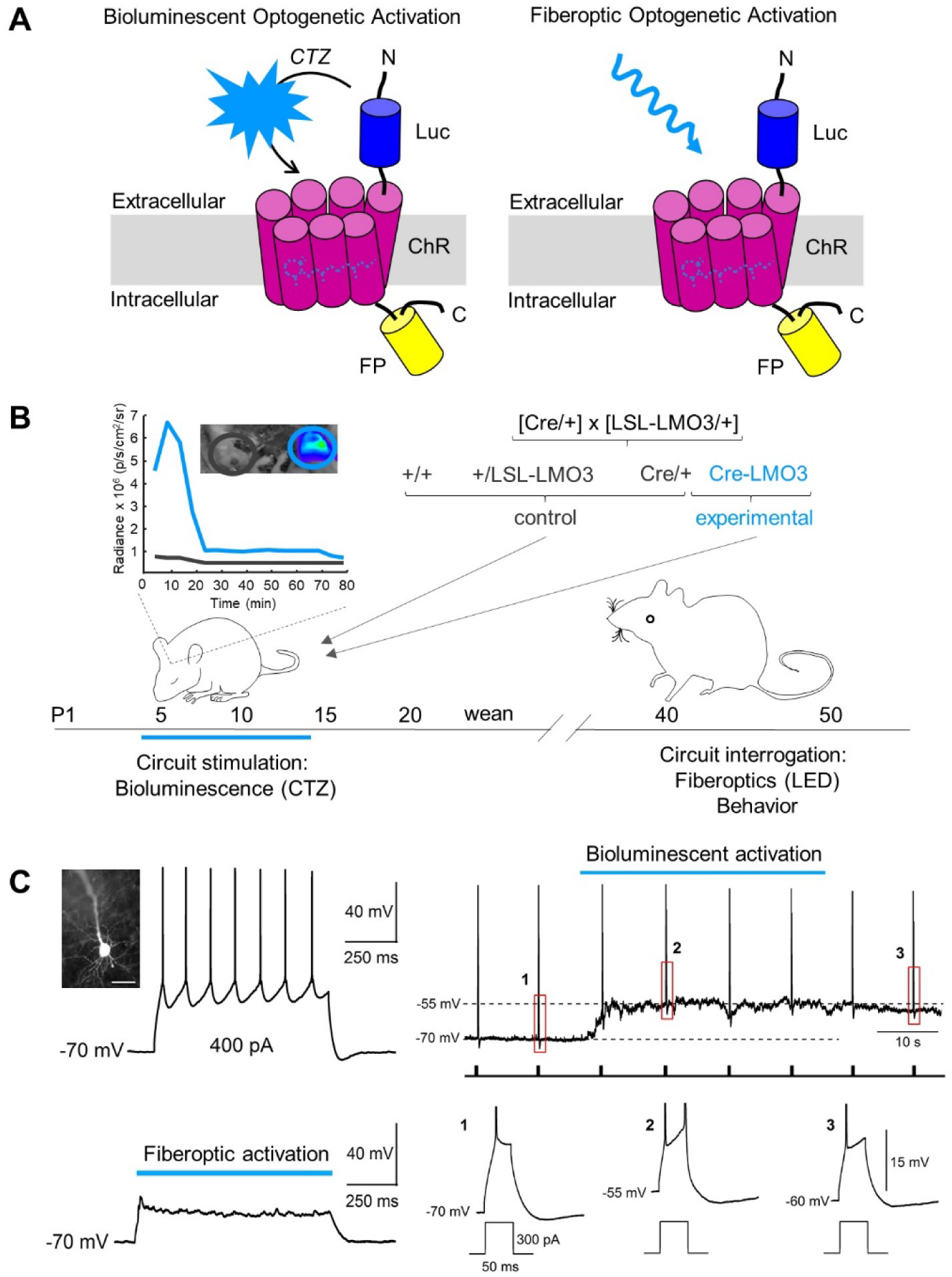
Bioluminescent optogenetics allows developmental chemogenetic drive of distinct neural populations for adult optogenetic circuit interrogation and behavioral testing. (A) Schematics of a luminopsin. A luciferase (Luc) is tethered to a channelrhodopsin (ChR); in addition, a fluorescent protein (FP) is fused to the C-terminus of the ChR as an expression marker. Application of the small molecule substrate coelenterazine (CTZ) results in production of photons and bioluminescent optogenetic activation of the nearby opsin (left). The same molecule is accessible to stimulation by a physical light source for standard fiberoptic optogenetic activation (right). (B) Experimental Design. Heterozygous Cre driver mice (Cre/+) were mated with heterozygous conditional (lox-stop-lox) luminopsin-3 mice (LSL-LMO3/+), generating three groups of control mice and one group of experimental mice expressing LMO3 in cells specified by the Cre driver (Cre-LMO3). All pups of a litter were injected once a day intraperitoneally with CTZ or vehicle postnatal days 4-14. Inset shows representative example of IVIS imaging of littermates from an Emx1-Cre/+ x LSL-LMO3/+ breeding. Pseudo-colored images show bioluminescence emitting only from the head of the Emx1-LMO3 double positive pup. ROIs plotted over time show bioluminescence in expressing mice to peak over the first 20 minutes after injection and to decline over the next hour. All groups were tested behaviorally starting at 2 months of age. Selected sets of Cre-LMO3 mice were used for detailed electrophysiological experiments, taking advantage of fiberoptic optogenetic interrogation. See also Figure S1. (C) Effect of CTZ on a prefrontal cortex layer 5 pyramidal cell expressing LMO3 at postnatal day 7. LMO3-expression and pyramidal cell identity was confirmed through suprathreshold square current injection (800 ms, 400 pA) together with biocytin staining (top left, scale bar: 50 µm) and 470 nm light illumination (1 second, lower left). Brief bath application of CTZ (300 µM) depolarized the cell and increased firing output to current pulse stimulation (300 pA, 50 ms, 0.1 Hz) during continuous membrane potential recording at -70 mV (right).

We induced hyperexcitation in developing mouse pups once a day during post-natal days 4-14, capturing the window of significant outgrowth and development of key cortical circuits. BL-OG has been used to activate or silence neurons in adult mice and rats, using various routes of administration of CTZ (Tung, Gutekunst and Gross, 2015; Berglund *et al.*, 2016; Park *et al.*, 2017; Gomez-Ramirez *et al.*, 2019; Yu *et al.*, 2019). To test the efficacy of BL-OG in pups, we tested for bioluminescence emission after intraperitoneal CTZ administration. Injection of CTZ resulted in bioluminescence emission within the first 10 – 15 minutes and declined over the following hour in LMO3 expressing, but not in non-expressing pups (Figure 1B - inset). To test the effect of CTZ on early postnatal neurons we recorded from LMO3-expressing layer 5 prefrontal cortex pyramidal neurons in slices from P7 Emx1-LMO3 pups (Figure 1C). In these functionally immature cells, we found that action potential output required large amplitude current injections and exposure to 470 nm light typically evoked only subthreshold depolarization. The ability to drive excitation of young pyramidal neurons with BL-OG was tested during continuous membrane potential recording (- 70 mV) while delivering periodic square current injections to evoke firing (0.1 Hz, 300 pA, 50 ms). Brief bath application of CTZ (300 µM) depolarized the membrane potential and increased the firing response to current stimulation (Figure 1C – lower right). Taken together, our results demonstrate the suitability of using a conditional BL-OG system for exploring the effects of developmental hyperexcitation.

### Developmental Hyperexcitation of Emx1 Pyramidal Neurons is Distinctive in Causing ASD Phenotypes

Developing pups were exposed to hyperexcitation of cortical pyramidal neurons directly (Emx1) or of populations affecting pyramidal neuron activity directly or indirectly (Pvalb, Dlx5/6, DAT; Figure 2A). Stimulation of selected neural populations was limited to the postnatal period P4-14, after which the animals received no further CTZ application. To test specifically whether pyramidal cell hyperactivity induces ASD symptoms and to test the general prediction that behavioral phenotypes will emerge from over-excitation of distinct neuronal populations during development, we assayed selected behaviors associated with neurodevelopmental disorders, including social interaction, stereotypic behavior, and anxiety (Figure 2B). Following selective activation of pyramidal neurons but not of interneurons or dopaminergic systems with BL-OG early in development generated the key ASD phenotypes of reduced social behavior and increased compulsivity, as well as hypo-locomotion. Developmentally hyperexcited Emx1-LMO3 mice show significantly reduced time interacting with the interaction mouse during the social approach test by nearly 50% compared to non-expressing littermates (students t-test: t(18) = 3.07, p = 0.0067, N = 9-11 per group; Figures 2B – Social Approach and S3A). Similarly, Emx1-LMO3 mice display over 40% reduced time spent interacting with the novel mouse in the social novelty test compared to non-expressing littermates; however, these differences do not reach statistical significance (students t-test: t(18) = 1.97, p = 0.0650; Figures 2B – Social Novelty and S3B). Pvalb-LMO3 mice demonstrate over 50% reduced time spent interacting with the novel mouse compared to non-expressing littermates (students t-test: t(21) = 2.19, p = 0.0398, N = 8-15 per group). Emx1-LMO3 mice were the only group to exhibit evidence of repetitive behaviors. Mice were observed for grooming behaviors in an open chamber for 10 minutes. Emx1-LMO3 mice show significantly increased time spent grooming by 250% compared to non-expressing littermates (student’s t-test: t(18) = -2.92, p = 0.0091, N = 9-11 per group; Figures 2B - Grooming and S3C). Emx1-LMO3 mice again were the only cohort to demonstrate altered exploration in open field. These mice show significantly reduced movements over time in open field (Two-way repeated measures ANOVA - Main effect between groups: F(1,14) = 11.56, p = 0.0043, N = 9-11 per group; Figures 2B – Open Field and S3D). All mice (expressing and non-expressing) show significant decline in movements over time, and Emx1-LMO3 show similar declines compared to non-expressing littermates (Main effect for Time: F(11,154) = 31.30, p < 0.0001; Interaction effect of Time x Group: F(11,154) = 1.71, p = 0.0764). When tested over 4 consecutive days, these results are maintained (Main effect between groups: F(1,14) = 22.58, p = 0.0003; Main effect for Time: F(3,42) = 7.77, p = 0.0003; Figure S3D). These behavioral effects were selective to pyramidal activation: Cortical cell activation by itself, in the form of interneuron drive, for example in the Dlx5/6 group, had opponent effects. Furthermore, the pyramidal activation was selective for ASD behavioral effects: developmental hyperexcitation of Emx1 neurons did not result in changes in anxiety for which imbalances in excitatory glutamate and inhibitory GABA circuits have been implicated (Depino, Tsetsenis and Gross, 2008). Indeed, developmental hyperexcitation of GABAergic Dlx5/6 neurons caused mice to display evidence of increased anxiety. These mice spend significantly reduced time on the open arms during the elevated plus maze by over 75% compared to non-expressing littermates (log-transformed data, student’s t-test: t(23) = 2.52, p = 0.0096, N = 8-22 per group; Figure 2B – Elevated Plus and Figure S3E). To ensure that none of the mouse lines were affected in basic coordination we employed a commonly used metric of basic learning and coordination, the rotarod. There were no significant differences on time to fall in any of the developmentally treated mice compared to non-expressing littermates (Figure 2B - Rotarod).

**Figure 2.**
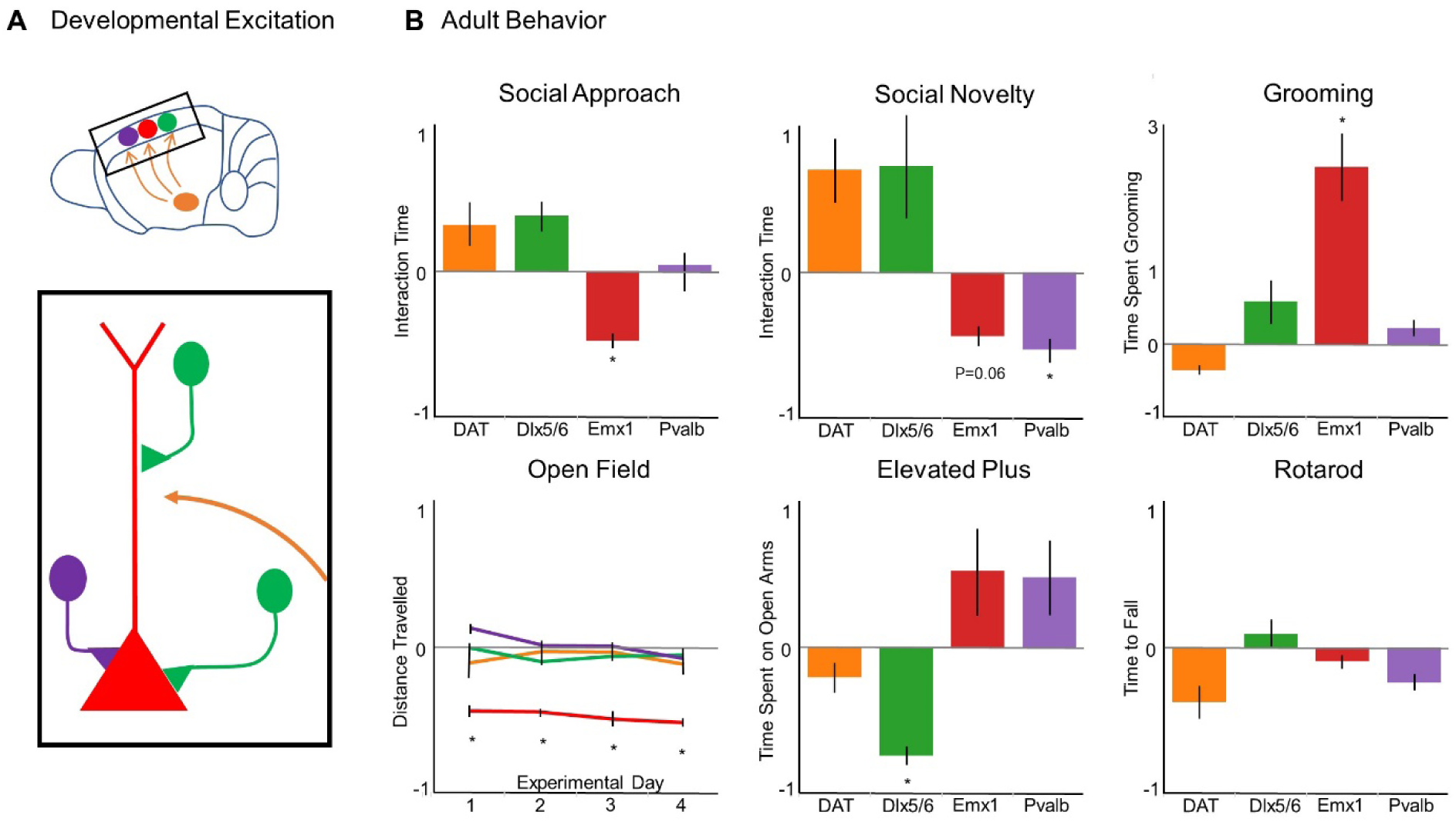
Developmental Hyperexcitation of Emx1 Pyramidal Neurons is Distinctive in Causing ASD Phenotypes. (A) Schematics of circuits targeted for developmental hyperexcitation. Color codes are used consistently for A and B: red – Emx1, green – Dlx5/6, purple – Pvalb, orange – DAT. (B) Adult behavior of developmentally hyperexcited mice. Each group of Cre (DAT, Dlx5/6, Emx1, Pvalb)-LMO3 mice is normalized to their non-LMO3 expressing controls. See also Figure S2. Social Approach/Interaction Test: Emx1-LMO3 mice show significantly reduced interaction time with another mouse. Social Novelty Test: Pvalb-LMO3 mice show significantly reduced interaction time with a second, novel mouse; Emx1-LMO3 mice show non-significantly reduced interaction (p = 0.065). Grooming Behaviors: Emx1-LMO3 mice show significantly elevated grooming behaviors. Open Field Test: Emx1-LMO3 mice show significantly reduced movements over 4 consecutive days of testing. Elevated Plus Maze: Dlx5/6-LMO3 mice show significantly reduced time spent on the open arms. Rotarod Test: None of the 4 animals groups show any significant differences on the accelerating rotarod compared to non-expressing controls. \**p*<.*05.*

### Developmental Pyramidal Hyperexcitation Leads to Decreased Cortico-Striatal Communication

Studies in humans and mouse models directly indicate that cortico-striatal communication is altered in neurodevelopmental disorders with basal ganglia association, such as Tourette’s and obsessive-compulsive disorders. Also, many autism models show altered connectivity between the cortex and the striatum (Peca *et al.*, 2011; Wang *et al.*, 2016; Morris, Baek and Voon, 2017; Nagarajan *et al.*, 2017; Abbott *et al.*, 2018; Martella *et al.*, 2018). To directly assay cortico-striatal communication in the relay of endogenous and controlled neocortical events, we measured the impact of naturally occurring neocortical bursts and induced optogenetic bursts on striatal evoked potential responses. Recordings were carried out in adult Emx1-LMO3 mice that had been treated during postnatal days 4-14 with CTZ (CTZ_P4-14_ mice) or with vehicle (VEH_P4-14_ mice). The bimodal feature of the LMOs enabled optogenetic access to Emx1 pyramidal neurons in both groups, and in CTZ_P4-14_ mice allowed optogenetic activation of the same neurons initially hyperexcited chemogenetically during development. We placed laminar electrodes in the medial prefrontal cortex and the dorsal striatum (Figure 3A). After 20 minutes of baseline, mice received acute stimulation to the prefrontal cortex through light pulses of 10s separated by 1-minute intervals, and recordings were continued for an hour. To identify changes in striatal responses, local field potentials (LFP) were captured after bursts of cortical activity to identify differences between baseline and light stimulus periods (Figure 3B-E). CTZ_P4-14_ mice show significantly reduced amplitude during baseline conditions (ANOVA: F(3,11752) = 51.28, p < 0.0001; Bonferroni post-hoc: p < .001; Figure 3B,C), suggesting weaker responses in the striatum to incoming stimulus. Similarly, CTZ_P4-14_ mice show no significant changes in amplitude during light stimulus (Bonferroni post-hoc: p = 0.913; Figure 3D). VEH_P4-14_ mice, by contrast, show a significant increase in amplitude during light stimulation (Bonferroni post-hoc: p < 0.001; Figure 3D). The frequency of LFPs during baseline shows no significant differences between groups; however, CTZ_P4-14_ mice show a decline in frequency during the light stimulus, while the VEH_P4-14_ group shows an increase (Mann-Whitney: z -1.964, p = 0.0496; Figure 3E). Based on the reduced amplitude of the LFPs, we conducted Fourier Transformations on the data to assess the power spectra within the striatum. CTZ_P4-14_ mice show significantly reduced power in the striatum in lower to mid frequency ranges including Theta, Alpha, and Beta ranges (Theta: F(3,108) = 3.19, p = 0.0265; Alpha: F(3,108) = 3.48, p = 0.0184; Beta: F(3,108) = 3.40, p = 0.0204; Figure 3F). The decrease in coupling in the beta and gamma bands is particularly notable given the robust *increase* in their expression during optogenetic drive of cortical pyramidal neurons (see Figure 4), reinforcing the conclusion of a functional decrease in connection between these key forebrain structures. Comparing the power spectra between baseline and stimulus conditions reveals no significant change in power among CTZ_P4-14_ mice during light stimulus in either the Theta, Alpha or Beta frequency ranges, indicating no response to increased incoming stimulus (Bonferroni post-hoc: p = 1.0; Figure 3G-I). VEH_P4-14_ mice, by contrast, show significantly elevated power during light stimulus compared to baseline in all 3 frequency ranges (Bonferroni post-hoc, Theta: p = 0.050; Alpha: p = 0.036; Beta: p = 0.042). Based on the increase in power and amplitude changes during light stimulation in VEH_P4-14_ mice, we assessed coherence between the cortex and striatum before, during and after light stimulation. CTZ_P4-14_ mice show significantly reduced coherence before and after light stimulus (Interaction of Time x Group: F(20,1128) = 1.87, p = 0.0117; Figure 3J), consistent with a broad decrease in coupling efficiency between the two structures. Coherence in both groups declines during light stimulus.

**Figure 3.**
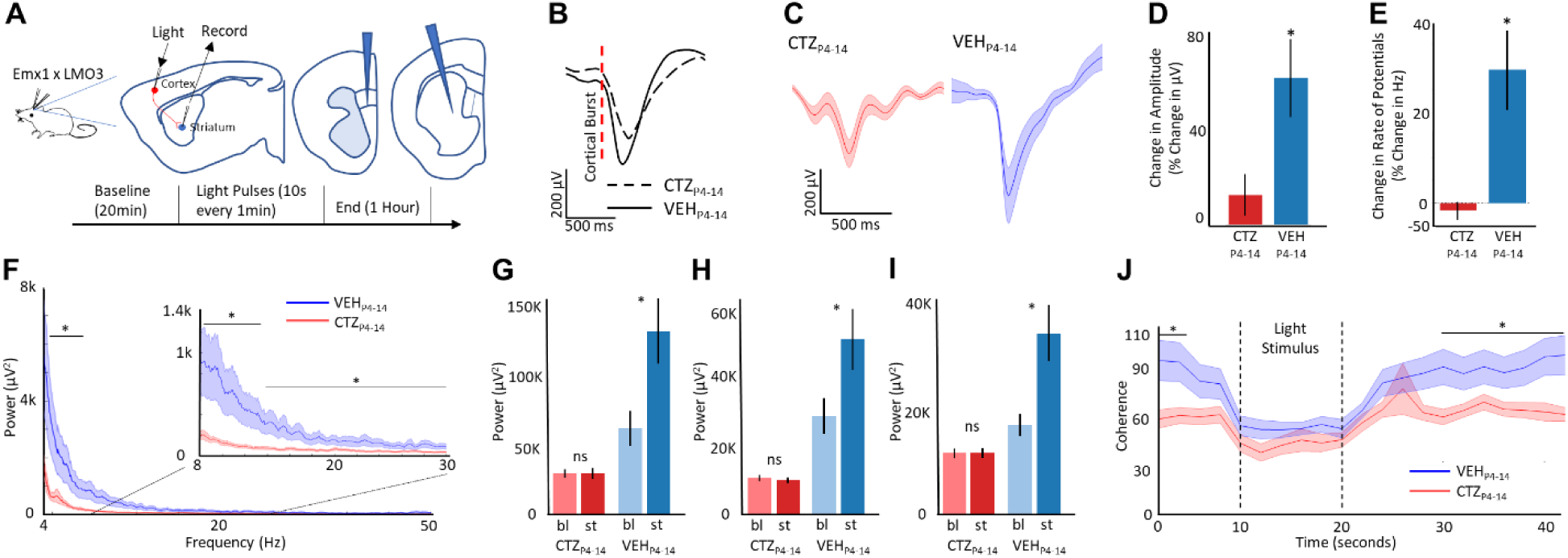
Developmental Pyramidal Hyperexcitation Leads to Decreased Cortico-Striatal Communication. (A) Schematic of experimental setup. Laminar probes were inserted in the prelimbic area of the medial prefrontal cortex and the striatum. (B) Striatal event-related Local Field Potentials (erLFPs) time locked to cortical action potential bursts (2 spikes in 10 ms) for adult Emx1-LMO3 mice developmentally stimulated (CTZ_P4-14_) and controls (VEH_P4-14_). (C) erLFPs during optogenetic stimulation: CTZ_P4-14_ mice (*red*) demonstrate smaller negative deflections compared to VEH_P4-14_ mice (*blue*). (D) Amplitude changes of erLFPs during optogenetic stimulus. Bars represent mean +/- SEM (E) Frequency changes among erLFPs during optogenetic stimulus. Bars represent mean +/- SEM (F) Power spectra of striatal neurons during light stimulus for CTZ_P4-14_ (*red*)- and VEH_P4-14_ (*blue*)- treated groups. (G-I) Average power spectra for the Theta (G), Alpha (H), or Beta (I) range during baseline (*bl*) and stimulus (*st*) conditions for both CTZ_P4-14_- and VEH_P4-14_- treated groups. Bars represent mean +/- SEM. (J) Coherence between cortex and striatum for both CTZ_P4-14_ (*red*)- and VEH_P4-14_ (*blue*)- treated mice before, during and after optogenetic stimulus. \**p*<.*05*

**Figure 4.**
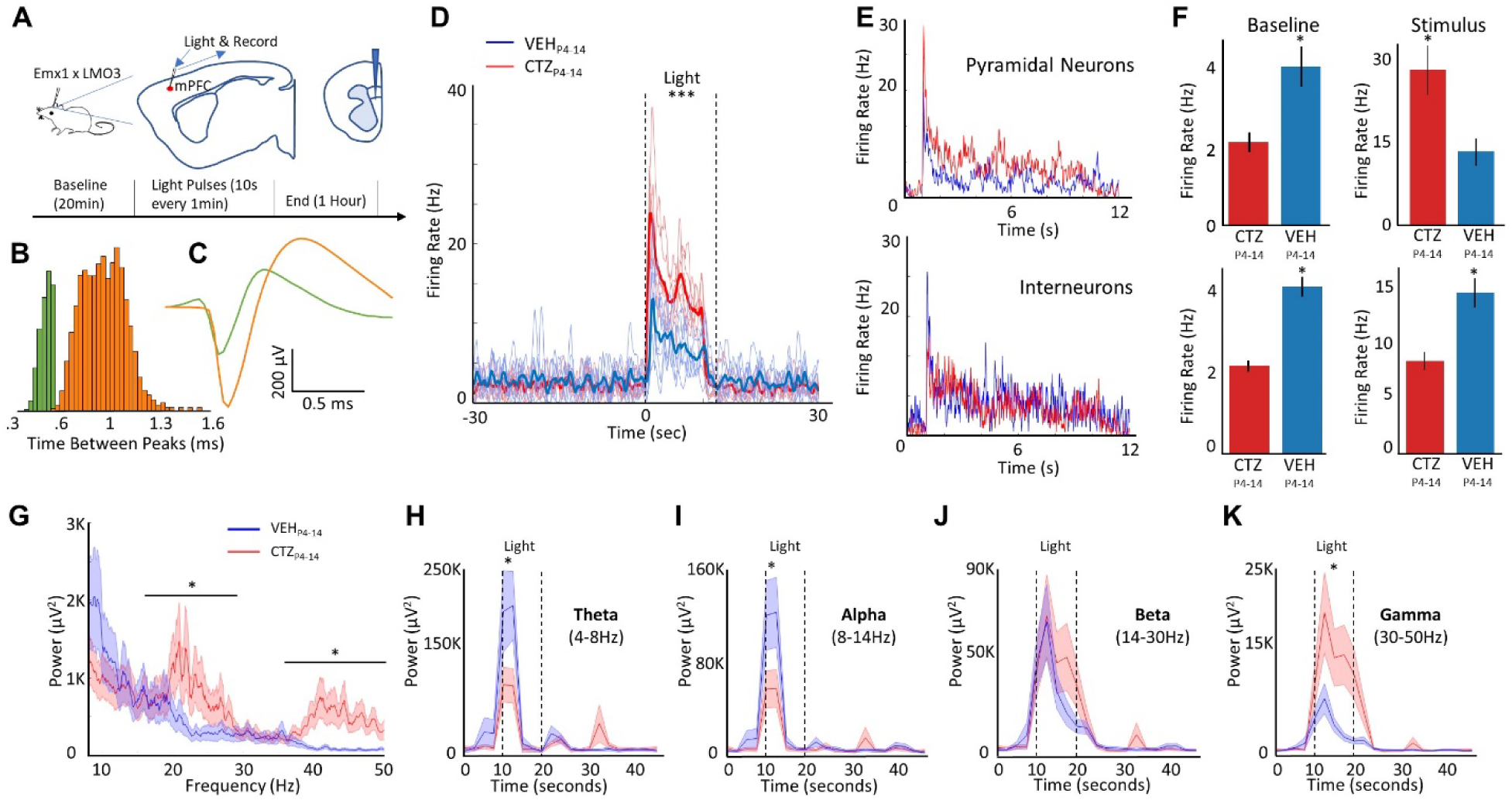
Developmental Pyramidal Hyperexcitation Leads to Decreased Baseline Firing, Enhanced Stimulus-Evoked Firing and Decreased Output Connectivity in Pyramidal Neurons. (A) Schematic of experimental setup. Laminar probes were inserted in the prelimbic area of the medial prefrontal cortex. (B) Waveforms were sorted based on time between peaks. Histogram of these peak to peak times shows a cluster of inhibitory neurons (*blue*) and pyramidal neurons (*orange*). (C) Traces of waveforms from putative inhibitory (*blue*) and pyramidal neurons (*orange*). (D) The group effects of optogenetic stimulation: CTZ_P4-14_-treated mice show a consistently larger response to stimulation compared to VEH_P4-14_-treated mice. (E) Light-dependent responses by pyramidal and interneuron populations: CTZ_P4-14_ mice (*red trace*) show far greater pyramidal neuron response to light stimulation (see upper panel), while VEH_P4-14_ mice (*blue trace*) show greater interneuron response to light stimulation (see lower panel). (F) Average firing rate for each group by neuron type and by timepoint (baseline or stimulus). Bars show mean ±SEM (upper panel – pyramidal neurons, lower panel – inhibitory neurons). (G) Power-spectra of cortical neurons during light stimulus. (H-K) Power over time before, during and after light stimulus for Theta (H), Alpha (I), Beta (J), and Gamma (K) frequency ranges. \**p*<.*05*

### Developmental Pyramidal Hyperexcitation Leads to Decreased Baseline Firing, Enhanced Stimulus-Evoked Firing and Decreased Output Connectivity in Pyramidal Neurons

Developmental hyperexcitation can be expected to be followed by homeostatic changes that would lead to reduced network-level excitability dynamics following sustained activity in specific neurons disproportionate to the rest of a network. At the single neuron level, predicted outcomes include reduced synaptic efficacy between the offending overexcited cell and its targets, and decreased excitability of these normal neighbors that receive the over-excited barrage. To directly test the impact of developmental hyperexcitation of pyramidal neurons, we interrogated neural and synaptic activity in neocortex of adult Emx1-LMO3 animals developmentally treated with CTZ (CTZ_P4-14_ mice) or with vehicle (VEH_P4-14_ mice). Recordings of cortical activity were conducted using a laminar probe traversing all layers of the cortex (Figure 4A). Continuous data was offline spike sorted using a high pass filtered at 250Hz. Spikes were identified as those that cross a threshold (−60µV). Spikes were separated using time between peaks to identify putative interneuron and pyramidal neuron subtypes (Figure 4B,C).

In agreement with the basic prediction of a tonic decrease in network-level excitability, we found decreased baseline firing in regular spiking neurons. However, direct optogenetic drive recruited a higher level of firing in these same cells. Across all mice and all trials, CTZ_P4-14_ mice, i.e. developmentally stimulated mice, show a consistently larger increase in firing rate, significantly greater than that of the VEH_P4-14_ mice, i.e. non-stimulated mice (Two-way Repeated Measures ANOVA, Main effect between groups: F(1,6) = 6.86, p = 0.0396; Figure 4D). No main effect was found for trial number, indicating the light stimulus did not produce differential effects regardless of when the trial occurred (Main effect for trial: F(6,36) = 1.38, p = 0.2494). No significant differences were found in the maximum firing rate between groups (Main effect between groups: F(1,6) = 1.30, p = 0.2977; data not shown), demonstrating that VEH_P4-14_ mice do reach a similar firing rate later in the light stimulation. This delay in maximal firing suggests some different excitatory/inhibitory dynamics between groups. Furthermore, the average firing rate after acute light stimulus is decreased in interneurons and increased in pyramidal neurons of CTZ_P4-14_ mice relative to VEH_P4-14_ mice (Figure 4E). This inverse relationship of baseline and stimulus-induced firing rate between pyramidal and interneurons is summarized in Figure 4F. During baseline, CTZ_P4-14_ mice show significantly reduced activity in the cortex compared to VEH_P4-14_ mice (Students T-test: t(7) = -6.082, p = 0.0005; Figure 4F). During light stimulus, by contrast, CTZ_P4-14_ mice show significantly *increased* firing rate compared to VEH_P4-14_ mice (t(7) = 2.541, p = 0.0386). Similarly, CTZ_P4-14_ mice show significantly reduced interneuron activity during baseline (Students T-test: t(7) = =2.526, p = 0.0395); however, CTZ_P4-14_ mice continue to show significantly reduced interneuron activity during light stimulation (Students T-test: t(7) = -2.41, p = 0.0468). The combination of these two findings suggests decreased excitatory synaptic efficacy from pyramidal neurons to neighboring targets is a key driver of reduced baseline activity. Specifically, sustained increased responses in pyramidal cells to optogenetic drive may reflect decreased efficacy of pyramidal input to interneurons, therefore reducing recruited di-synaptic inhibition when pyramidal neurons are driven.

Studies have demonstrated that altered excitatory-inhibitory dynamics would result in altered neuronal oscillations (Brunel and Wang, 2003; Atallah and Scanziani, 2009). These studies suggest that an altered ratio towards excitation would result in a shift in neural oscillations to higher power in the lower Gamma range (30-80Hz). To assess whether these dynamics are altered in CTZ_P4-14_ mice, we conducted fast Fourier transforms to obtain power spectra by frequency (Figure 4G). During light stimulus, CTZ_P4-14_ mice show significantly increased power in the Beta range (Student’s T-test: t(6) = 2.517, p = 0.0455), and the Gamma range (t(6) = 2.732, p = 0.0341), suggesting these mice may experience altered excitation/inhibition balance. Given the elevated power in the Beta and Gamma range among CTZ_P4-14_ mice, we assessed how these oscillations change over time. Fast Fourier Transforms were performed before, during, and after light stimulation to assess the change in power over time. VEH_P4-14_ mice and CTZ_P4-14_ mice exhibit different responses to light stimulation. In the lower frequency ranges such as Theta and Alpha, CTZ_P4-14_ mice show significantly reduced power in response to light stimulation (Two-way repeated measures – Interaction of time x group: Theta – F(20,1128) = 3.21, p < 0.0001; Alpha – F(20,1128) = 3.49, p < 0.0001; Figure 4H,I). The differences occur within the first few seconds of light stimulation (Bonferroni post-hoc: Theta – p = 0.0203; Alpha – p = 0.0302), with the power returning to baseline levels roughly half-way through the light stimulus. No significant differences over time were noted in the Beta frequency range (F(20,1128) = 0.78, p = 0.7392; Figure 4J). In the higher ranges, such as Gamma, CTZ_P4-14_ mice show significantly higher power compared to VEH_P4-14_ mice (Two-way repeated measures – Interaction of time x group: F(20,1128) = 5.40, p < 0.0001; Figure 4K). In both the Beta and Gamma range, CTZ_P4-14_ mice show a longer increase in power, lasting the entire length of the light stimulation (Figure 4J,K).

## Discussion

Here, we tested in healthy animals the impact of early enhancement in neural activity within specific cell types. We found that early hyperexcitation alone for a limited time window is sufficient to create irreversible adult neurophysiological and behavioral phenotypes, in the absence of any other genetic or environmental insults. Moreover, early postnatal interference with neural activity through hyperexcitation of distinct populations resulted in distinct behavioral symptoms, indicating that this is not a non-specific effect. In addition to causally linking developmental neural activity with adult behavior, our results provide insights into the dynamics of neural networks responding to aberrant activities during development.

Our findings directly indicate that brief and highly selective alterations in activity in neocortical pyramidal neurons may be a key common denominator in ASD. As such, they provide a potential causal explanation for ASD phenotypes being correlated with environmental toxins, infectious agents, stress, and a multitude of genetic and epigenetic origins.

The methodological use of BL-OG here uniquely enabled this work. Combined chemogenetic and optogenetic access to the same molecule minimized genetic alteration load as we only had to employ one actuator. This strategy provides a potential platform to identify novel therapeutic approaches addressing the etiologies of distinct behavioral hallmarks irrespective of the heterogeneous affronts converging on them.

The experimental system introduced here will allow dissection of precise timing of activity imbalance, interrogation of neural subpopulations, and assessment of an expanded repertoire of behavioral tests, thus providing opportunities to identify vulnerable periods during development and provide key behavioral symptoms of neurodevelopmental disorders with structural and functional correlates. Our approach offers an experimental framework for delineating developmental network disturbances and their consequences in establishing adult neural connections and their resulting behaviors in the general field of neurodevelopmental disorders.

## EXPERIMENTAL MODEL AND SUBJECT DETAILS

### Animals

All experiments involving animals were carried out following the guidelines and protocols approved by the Institutional Animal Care and Use Committee at Central Michigan University and were in compliance with the US National Research Council’s Guide for the Care and Use of Laboratory Animals, the US Public Health Service’s Policy on Humane Care and Use of Laboratory Animals, and Guide for the Care and Use of Laboratory Animals.

Mice were group-housed in ventilated cages under 12-hour reverse light cycle, provided with tap water and standard chow and allowed to feed *ad libitum*.

Experimental animals were generated by crossing LSL-LMO3 mice (see below) with the following Cre driver lines: Emx1-Cre, JAX# 005628; Pvalb-Cre, JAX# 017320; Dlx5/6-Cre, JAX# 008199; DAT-Cre, JAX# 006660.

## METHOD DETAILS

### Generation of LSL-LMO3 mice

A ROSA26 targeting construct placing LMO3 (sbGLuc-VChR1-EYFP; Berglund *et al.*, 2016) under conditional (lox-stop-lox, LSL) control of the strong ubiquitous CAG promoter was generated by replacing the tdTomato gene in the Allen Brain Institute’s Ai9 targeting vector (CAG-floxed tdTomato; Addgene plasmid 22799; contributed by Hongkui Zeng; Madisen et al. 2010). Embryonic stem cells were homologously targeted via electroporation of (129×1/SvJ x 129S1/Sv)F1-*Kitl*^*+*^-derived R1 embryonic stem (ES) cells (Nagy *et al.*, 1993), injected into blastocysts, and male chimeras were crossed with C57BL/6 females. Heterozygous LSL-LMO3 mice were further crossed to C57BL/6. The PGK-neo marker is flanked by a pair of PhiC31 recognition sites (*attB*/*attP*) and was deleted from the LSL-LMO3 line by crossing with PhiC31 deleter mice (Stock #007743, Jackson Laboratory; kindly provided by Dr. Hongkui Zeng, Allen Brain Institute). Routine genotyping for detecting the presence of the conditional alleles was done using forward primer 5’-ATGTCTGGATCCCCATCAAG, and reverse primer 5’-TCCGAAGCCAACCTTCACAGTAAC (Zhu *et al.*, 2016).

### IVIS Imaging

Mouse pups aged post-natal day 4 were injected with CTZ intraperitoneally at a dose of 10µg/g. Pups were anesthetized on ice prior to being placed in the IVIS chamber. Images were taken in 5-minute bins over a 1-hour period and quantified using radiance (Perkin-Elmer, IVIS Lumina LT, Living Image Software).

### Ex vivo recording

Acute brain slices were prepared from P7 Emx1-LMO3 pups. Briefly, mice were anaesthetized via inhalation of isoflurane and, following decapitation, the brain was isolated and placed in ice-cold cutting solution containing (in mM): 92 NMDG, 2.5 KCl, 0.5 CaCl2, 10 MgSO4, 30 NaHCO3, 1.25 NaH2PO4, 20 HEPES, 2 Thiourea, 5 Na-ascorbate, 3 Na-pyruvate and 25 D-glucose (310 mOsm/kg, pH 7.3-7.4). Coronal slices (300 *μ*m) from prefrontal cortex were cut using a Leica VT1000s vibratome and stored in a recovery solution containing (in mM): 92 NaCl, 2.5 KCl, 2 CaCl2, 2 MgSO4, 30 NaHCO3, 1.25 NaH2PO4, 20 HEPES, 2 Thiourea, 5 Na-ascorbate, 3 Na-pyruvate and 25 D-glucose (310 mOsm/kg, pH 7.3-7.4). After a one hour recovery period, a slice was transferred to a recording chamber mounted on an upright microscope (Olympus BX51WI) and perfused with aCSF containing (in mM): 121 NaCl, 2.8 KCl, 1 NaH2PO4, 26 NaHCO3, 2 CaCl2, 2 MgCl2 and 15 D-glucose (300 mOsm/kg, pH 7.3-7.4) at a rate of 4 ml/min. All solutions were bubbled with a gas mixture of 95% O2 and 5% CO2. Whole-cell current clamp recordings were made using a Multiclamp 700b amplifier, a Digidata 1440 digitizer together with pClamp software (Molecular Devices). Borosilicate glass micropipettes were manufactured using a PC-100 puller (Narishige) and had resistances of 4–6 M*Ω*. Pipettes were filled with intracellular solution containing (in mM): 130 K-gluconate, 10 KCl, 15 HEPES, 5 Na2-phosphocreatine, 4 Mg-ATP, 0.3 Na-GTP and 0.5% biocytin (300 mOsm/kg, pH 7.3). LMO3-expressing L5 pyramidal neurons were visually targeted using epifluorescence microscopy together with a CMOS camera (Hamamatsu OCRA Fusion). The membrane potential response to current injection, 470 nm light and CTZ was recorded at -70 mV. Data was sampled at 10 kHz, filtered at 3 kHz and analyzed in Igor Pro (WaveMetrics). Access resistance was <15 M*Ω* at start of recordings. Cells displaying a change in access resistance >15% over the course of the experiment were discarded. The liquid junction potential was not corrected for.

### Microscopy

Immunofluorescent images were captured on a Zeiss AxioCam M2 microscope using a 20x objective and digitized using the ZEN software (Carl Zeiss Inc., Thornwood, NY, USA).

### Treatment of pups

The entire litter from heterozygous breeding pairs of Cre-mice and LSL-LMO3 received CTZ (water soluble native coelenterazine; Prolume Inc., cat# 3031) or vehicle (water soluble carrier without CTZ; Prolume Inc., cat# 3031C). CTZ or vehicle were injected intraperitoneally at a dose or volume equivalent of 10µg/g once per day during p4-14. Mice were then weaned, genotyped, and group-housed until used for behavioral and recording experiments.

### Behavioral tests

All behavioral tests were performed with age-matched male and female littermates, starting at 2 – 3 months old and continuing over a 3-4 months period. We did not observe gender differences and thus male and female mice were pooled, generating groups of 8 – 22 animals. Mice were moved between holding room and behavioral suite, located within the same facility. Behavioral tests were carried out during the day in rooms under reverse light cycle. Testing was carried out by individuals blinded to experimental conditions. Scoring for each test was done by 2 independent individuals blinded to experimental conditions.

#### Three Chamber Test

Social behavior was tested using the 3-chamber test (Crawley, 2007; Yang, Silverman and Crawley, 2011). Animals were placed in a 27” × 14” chamber with 3 segments of equal size (Medendorp *et al.*, 2018). Animals were allowed to explore the arena for 5 min to habituate to the apparatus. Mice were tested first for social approach, which has been demonstrated to relate to social deficits found in autism (Yang, Silverman and Crawley, 2011). Mice were additionally tested for social novelty immediately following the social approach test.

##### Social Approach

Two identical plastic cylinders were placed in either of the external chambers. These cylinders were clear, with regular holes to allow for visual and olfactory stimuli. A sex-matched, non-familiar mouse was placed in one of these cylinders. Experimental mice were placed in the middle, empty section of the 3-chamber apparatus and allowed to roam for 5 minutes. Social interaction was characterized by time spent interacting with the stationary mouse, i.e. time spent within a 1-inch radius of the mouse compared to within a 1-inch radius of the empty cylinder.

##### Social Novelty

Following the social approach test, a novel, sex-matched, non-familiar mouse was placed in the previously empty cylinder. Experimental mice were again allowed to roam for 5 minutes. Social behavior was characterized by time spent interacting with the novel mouse compared to the previous mouse. The chamber was cleaned with 70% ethanol between testing mice.

#### Elevated Plus Maze

Mice were placed in the center of an elevated, plus-shaped apparatus (30.5” × 30.5” arms, 30.5” from the ground). Two of the external arms were covered, and 2 were open. Mice were allowed to roam the apparatus for 5 minutes. Mice were assessed for time spent on the open arms, with more time spent on the open arms indicative of less anxiety.

#### Open Field

Experimental mice were placed in a 17” × 9” cage and allowed to roam for 60 minutes. Movements were tracked by laser grid and analyzed using Hamilton-Kinder™ motor monitor software. Tests were repeated over 4 consecutive days using the same cage. Overall ambulation of the mice was assessed to establish normal exploratory behavior (Crawley, 1985).

#### Grooming Observation

Mice were observed in a 17” × 9” cage for a period of 10 minutes. Periods of grooming were noted and totaled over the 10-minute period.

#### Rotarod

Mice were placed on an accelerating rotarod and timed until they were unable to remain on the spinning rod. Mice were trained on the rotarod every day for 4 days prior to testing. The rotarod accelerated from 10-40 revolutions per minute (rpm) over the course of 30 seconds. At the end of the 30 seconds the rotarod remained at 40 rpm until 180 seconds or until mice were unable to stay on the rotarod. Mice were tested for 3 trials of the rotarod and the scores averaged.

### In vivo recording

Mice were anesthetized with urethane at 1.5g/kg and mounted on a stereotax (Kopf Instruments). Laminar optoelectrodes were inserted in the prelimbic area of the medial prefrontal cortex (mPFC; 2mm anterior to bregma, 0.4mm lateral, and 2mm ventral). The striatal electrode (20° lateral offset) was placed 0.7mm anterior to bregma, 3mm lateral, and 2.6mm ventral. Laminar probes consisted of single shank, 32-channel silicon probes with a fiber optic 50µm above the highest recording site (A1×32 Poly2-5mm-50s-177-OA32LP, Neuronexus Technologies). Data was sampled at 30kHz and passed through a digital amplifier (Cereplex-µ, Blackrock Microsystems), and directed through HDMI to the Cereplex Direct data acquisition box (Blackrock Microsystems).

VChR1 photostimulation was carried out using a PlexBright optogenetic stimulation system (Plexon Inc.) with a blue LED module (465 nm). Mice were recorded for 20 minutes to establish a baseline. At 20 minutes, blue light was applied through the fiber optic at 300µA intensity for 10 seconds. Each light pulse was separated by 1 minute. Recordings were allowed to continue 1 hour after injection to assess response to stimulation. After recordings, brains were extracted and sectioned to confirm electrode placement.

## QUANTIFICATION AND STATISTICAL ANALYSIS

All analysis was carried out using MATLAB and SPSS software. Data are displayed as mean±standard error of the mean (SEM). Behavior tests were tested using student’s t-test between expressing and nonexpressing mice. Open field was tested using two-way repeated measures ANOVAs.

Electrophysiological data was high pass filtered at 250Hz to extract spikes and low pass filtered at 300Hz to extract LFPs. Spike data was thresholded at -63µV and sorted for each channel based on waveform characteristics using Principal Components Analysis (PCA). Spikes were binned to calculate frequency of firing over time. Differences between groups were assessed using two-way repeated measures ANOVAs (repeat trials per mouse).

Fast Fourier transforms were carried out using MATLAB **fft()** function. LFP waveform data were converted from time to frequency, producing power spectral density histograms. Data was pulled from a 12-second interval spanning 1second prior to light stimulus and 1 second after to encompass the entire 10 second light stimulation period. This was repeated for each light trial. Data was quantified from various frequency ranges including Delta (0-4Hz), Beta (4-8Hz), Theta (8-14Hz), and Gamma (30-100Hz).

## SUPPLEMENTAL FIGURES

**Supplementary Figure 1.**
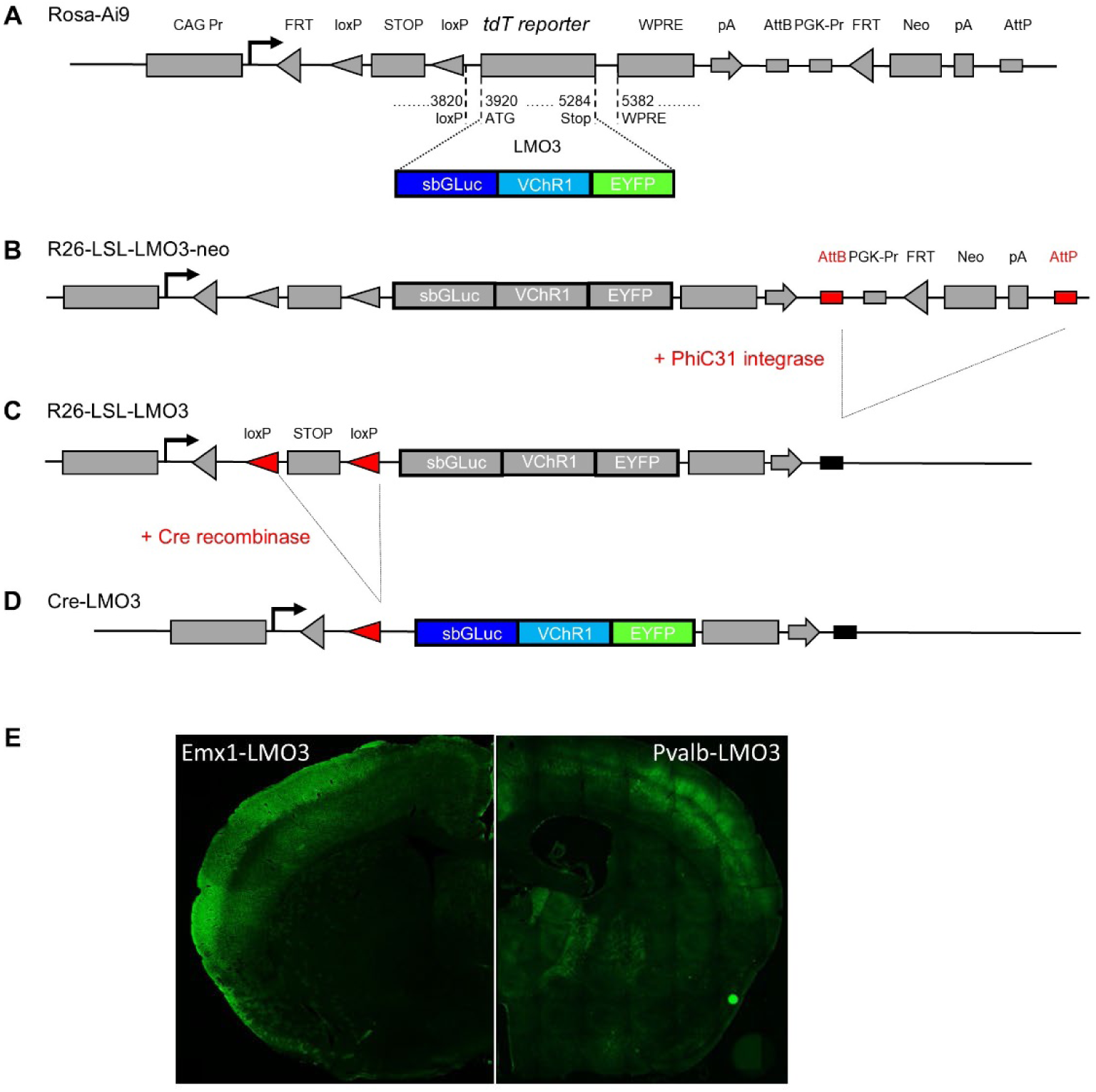
Conditional (lox-stop-lox) LMO3 mice. (A) Schematic of RosaAi9 targeting vector. The tdTomato reporter was replaced by the coding region for LMO3. (B) Schematic of R26-LSL-LMO3-neo construct used for generating LSL-LMO3-neo embryonic stem cells and germline transmitting LSL-LMO3-neo mice. (C) Schematic showing the modified Rosa26 locus after crossing LSL-LMO3-neo mice to PhiC31 integrase expressing mice, resulting in removal of the neo reporter in R26-LSL-LMO3 mice. These mice were crossed with the various Cre driver lines. (D) Schematic of Rosa26 locus in cells co-expressing Cre recombinase, resulting in removal of the Stop sequence and expression of LMO3 in Cre-LMO3 mice. (E) Fluorescence images of brain cross sections of Emx1-LMO3 (left) and Pvalb-LMO3 (right) mice.

**Supplementary Figure 2.**
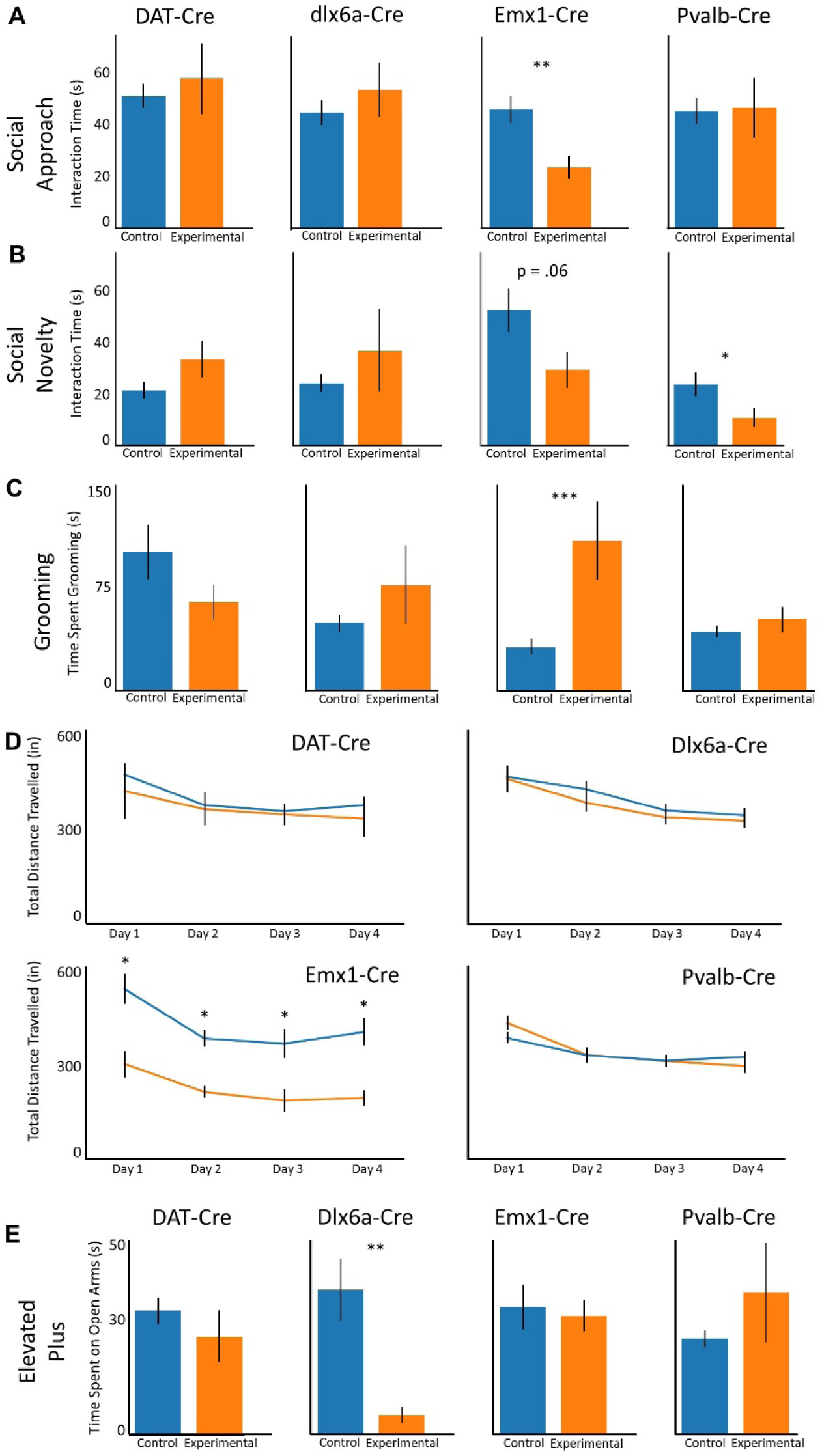
Behavioral tests. (A – E) Results of behavioral tests showing measurements for control (non-expressing) and experimental (LMO3-expressing) animals treated with CTZ during postnatal development (P4-14). Bars show mean ±SEM.

## References

Abbott, A. E. et al. (2018) ‘Repetitive behaviors in autism are linked to imbalance of corticostriatal connectivity: a functional connectivity MRI study.’, Social cognitive and affective neuroscience. England, 13(1), pp. 32–42. doi: 10.1093/scan/nsx129.

Antoine, M. W. et al. (2019) ‘Increased Excitation-Inhibition Ratio Stabilizes Synapse and Circuit Excitability in Four Autism Mouse Models.’, Neuron. United States, 101(4), pp. 648-661.e4. doi: 10.1016/j.neuron.2018.12.026.

Atallah, B. V. and Scanziani, M. (2009) ‘Instantaneous Modulation of Gamma Oscillation Frequency by Balancing Excitation with Inhibition’, Neuron, 62(4), pp. 566–577. doi: 10.1016/j.neuron.2009.04.027.

Backman, C. M. et al. (2006) ‘Characterization of a mouse strain expressing Cre recombinase from the 3’ untranslated region of the dopamine transporter locus.’, Genesis (New York, N.Y. : 2000). United States, 44(8), pp. 383–390. doi: 10.1002/dvg.20228.

Balsters, J. H., Mantini, D. and Wenderoth, N. (2018) ‘Connectivity-based parcellation reveals distinct cortico-striatal connectivity fingerprints in Autism Spectrum Disorder.’, NeuroImage. United States, 170, pp. 412–423. doi: 10.1016/j.neuroimage.2017.02.019.

Berglund, K. et al. (2013) ‘Light-Emitting Channelrhodopsins for Combined Optogenetic and Chemical-Genetic Control of Neurons’, PLoS ONE, 8(3), p. e59759.

Berglund, K. et al. (2016) ‘Luminopsins integrate opto- and chemogenetics by using physical and biological light sources for opsin activation’, Proceedings of the National Academy of Sciences of the United States of America, 113(3), pp. E358–E367. doi: 10.1073/pnas.1510899113.

Berglund, K. and Gross, R. E. (2019) ‘Opto-chemogenetics with luminopsins: A novel avenue for targeted control of neuronal activity.’, Journal of neuroscience research. United States. doi: 10.1002/jnr.24473.

Brunel, N. and Wang, X. J. (2003) ‘What determines the frequency of fast network oscillations with irregular neural discharges? I. Synaptic dynamics and excitation-inhibition balance’, Journal of Neurophysiology. American Physiological Society, 90(1), pp. 415–430. doi: 10.1152/jn.01095.2002.

Chaste, P. and Leboyer, M. (2012) ‘Autism risk factors: genes, environment, and gene-environment interactions.’, Dialogues in clinical neuroscience. France, 14(3), pp. 281–292.

Crawley, J. N. (1985) ‘Exploratory behavior models of anxiety in mice.’, Neuroscience and biobehavioral reviews. United States, 9(1), pp. 37–44. doi: 10.1016/0149-7634(85)90030-2.

Crawley, J. N. (2007) What’s Wrong With My Mouse?: Behavioral Phenotyping of Transgenic and Knockout Mice. John Wiley & Sons.

Depino, A. M., Tsetsenis, T. and Gross, C. (2008) ‘GABA homeostasis contributes to the developmental programming of anxiety-related behavior.’, Brain research. Netherlands, 1210, pp. 189–199. doi: 10.1016/j.brainres.2008.03.006.

Fuccillo, M. V (2016) ‘Striatal Circuits as a Common Node for Autism Pathophysiology.’, Frontiers in neuroscience. Switzerland, 10, p. 27. doi: 10.3389/fnins.2016.00027.

Geschwind, D. H. (2011) ‘Genetics of autism spectrum disorders.’, Trends in cognitive sciences. England, 15(9), pp. 409–416. doi: 10.1016/j.tics.2011.07.003.

Gomez-Ramirez, M. et al. (2019) ‘The BioLuminescent-OptoGenetic in vivo Response to Coelenterazine is Proportional, Sensitive and Specific in Neocortex’, J Neurosci Res. doi: 10.1101/709931.

Gorski, J. A. et al. (2002) ‘Cortical excitatory neurons and glia, but not GABAergic neurons, are produced in the Emx1-expressing lineage.’, The Journal of neuroscience : the official journal of the Society for Neuroscience, 22(15), pp. 6309–14. doi: 20026564.

Hippenmeyer, S. et al. (2005) ‘A developmental switch in the response of DRG neurons to ETS transcription factor signaling.’, PLoS biology. United States, 3(5), p. e159. doi: 10.1371/journal.pbio.0030159.

Kuo, H.-Y. and Liu, F.-C. (2017) ‘Valproic acid induces aberrant development of striatal compartments and corticostriatal pathways in a mouse model of autism spectrum disorder.’, FASEB journal : official publication of the Federation of American Societies for Experimental Biology. United States, 31(10), pp. 4458–4471. doi: 10.1096/fj.201700054R.

Lord, C. et al. (2000) ‘Autism spectrum disorders.’, Neuron. United States, 28(2), pp. 355–363. doi: 10.1016/s0896-6273(00)00115-x.

Madisen, L. et al. (2010) ‘A robust and high-throughput Cre reporting and characterization system for the whole mouse brain.’, Nature neuroscience, 13(1), pp. 133–40. doi: 10.1038/nn.2467.

Martella, G. et al. (2018) ‘The neurobiological bases of autism spectrum disorders: the R451C-neuroligin 3 mutation hampers the expression of long-term synaptic depression in the dorsal striatum.’, The European journal of neuroscience. France, 47(6), pp. 701–708. doi: 10.1111/ejn.13705.

Medendorp, W. E. et al. (2018) ‘Altered Behavior in Mice Socially Isolated During Adolescence Corresponds With Immature Dendritic Spine Morphology and Impaired Plasticity in the Prefrontal Cortex.’, Frontiers in behavioral neuroscience. Switzerland, 12, p. 87. doi: 10.3389/fnbeh.2018.00087.

Monory, K. et al. (2006) ‘The endocannabinoid system controls key epileptogenic circuits in the hippocampus.’, Neuron. United States, 51(4), pp. 455–466. doi: 10.1016/j.neuron.2006.07.006.

Moore, C. I. and Berglund, K. (2019) ‘BL-OG: BioLuminescent-OptoGenetics.’, Journal of neuroscience research. United States. doi: 10.1002/jnr.24575.

Morris, L. S., Baek, K. and Voon, V. (2017) ‘Distinct cortico-striatal connections with subthalamic nucleus underlie facets of compulsivity.’, Cortex; a journal devoted to the study of the nervous system and behavior, 88, pp. 143–150. doi: 10.1016/j.cortex.2016.12.018.

Nagarajan, N. et al. (2017) ‘Corticostriatal circuit defects in Hoxb8 mutant mice’, Molecular Psychiatry. Nature Publishing Group, 23(9). doi: 10.1038/mp.2017.180.

Nagy, A. et al. (1993) ‘Derivation of completely cell culture-derived mice from early-passage embryonic stem cells.’, Proceedings of the National Academy of Sciences of the United States of America, 90(18), pp. 8424–8. Available at: http://www.pubmedcentral.nih.gov/articlerender.fcgi?artid=47369&tool=pmcentrez&rendertype=abstract (Accessed: 20 December 2015).

Nicolini, C. and Fahnestock, M. (2018) ‘The valproic acid-induced rodent model of autism.’, Experimental neurology. United States, 299(Pt A), pp. 217–227. doi: 10.1016/j.expneurol.2017.04.017.

Park, S. Y. et al. (2017) ‘Novel luciferase-opsin combinations for improved luminopsins’, Journal of Neuroscience Research. doi: 10.1002/jnr.24152.

Peca, J. et al. (2011) ‘Shank3 mutant mice display autistic-like behaviours and striatal dysfunction.’, Nature. England, 472(7344), pp. 437–442. doi: 10.1038/nature09965.

Peixoto, R. T. et al. (2016) ‘Early hyperactivity and precocious maturation of corticostriatal circuits in Shank3B(-/-) mice’, Nature Neuroscience, 19(5), pp. 716–724. doi: 10.1038/nn.4260.

Peixoto, R. T. et al. (2019) ‘Abnormal Striatal Development Underlies the Early Onset of Behavioral Deficits in Shank3B(-/-) Mice.’, Cell reports. United States, 29(7), pp. 2016-2027.e4. doi: 10.1016/j.celrep.2019.10.021.

Ramaswami, G. and Geschwind, D. H. (2018) ‘Genetics of autism spectrum disorder.’, Handbook of clinical neurology. Netherlands, 147, pp. 321–329. doi: 10.1016/B978-0-444-63233-3.00021-X.

Rothwell, P. E. et al. (2014) ‘Autism-associated neuroligin-3 mutations commonly impair striatal circuits to boost repetitive behaviors.’, Cell. United States, 158(1), pp. 198–212. doi: 10.1016/j.cell.2014.04.045.

Roullet, F. I., Lai, J. K. Y. and Foster, J. A. (2013) ‘In utero exposure to valproic acid and autism--a current review of clinical and animal studies.’, Neurotoxicology and teratology. United States, 36, pp. 47–56. doi: 10.1016/j.ntt.2013.01.004.

Selimbeyoglu, A. et al. (2017) ‘Modulation of prefrontal cortex excitation/inhibition balance rescues social behavior in CNTNAP2-deficient mice.’, Science translational medicine. United States, 9(401). doi: 10.1126/scitranslmed.aah6733.

Shofty, B. et al. (2019) ‘Autism-associated Nf1 deficiency disrupts corticocortical and corticostriatal functional connectivity in human and mouse.’, Neurobiology of disease. United States, 130, p. 104479. doi: 10.1016/j.nbd.2019.104479.

Song, D. et al. (2018) ‘Manipulation of hippocampal CA3 firing via luminopsins modulates spatial and episodic short-term memory, especially working memory, but not long-term memory.’, Neurobiology of learning and memory. United States, 155, pp. 435–445. doi: 10.1016/j.nlm.2018.09.009.

Tung, J. K., Gutekunst, C.-A. and Gross, R. E. (2015) ‘Inhibitory luminopsins: genetically-encoded bioluminescent opsins for versatile, scalable, and hardware-independent optogenetic inhibition’, Scientific Reports, 5, p. 14366. doi: 10.1038/srep14366.

Uddin, L. Q., Supekar, K. and Menon, V. (2013) ‘Reconceptualizing functional brain connectivity in autism from a developmental perspective.’, Frontiers in human neuroscience. Switzerland, 7, p. 458. doi: 10.3389/fnhum.2013.00458.

Wang, X. et al. (2016) ‘Altered mGluR5-Homer scaffolds and corticostriatal connectivity in a Shank3 complete knockout model of autism’, Nature Communications, 7. doi: 10.1038/ncomms11459.

Watts, T. J. (2008) ‘The pathogenesis of autism.’, Clinical medicine. Pathology. New Zealand, 1, pp. 99–103. doi: 10.4137/cpath.s1143.

Yang, M., Silverman, J. L. and Crawley, J. N. (2011) ‘Automated three-chambered social approach task for mice’, Current Protocols in Neuroscience, Chapter 8, p. Unit 8.26. doi: 10.1002/0471142301.ns0826s56.

Yizhar, O. et al. (2011) ‘Neocortical excitation/inhibition balance in information processing and social dysfunction.’, Nature. England, 477(7363), pp. 171–178. doi: 10.1038/nature10360.

Yu, S. P. et al. (2019) ‘Optochemogenetic Stimulation of Transplanted iPS-NPCs Enhances Neuronal Repair and Functional Recovery after Ischemic Stroke.’, The Journal of neuroscience : the official journal of the Society for Neuroscience. United States, 39(33), pp. 6571–6594. doi: 10.1523/JNEUROSCI.2010-18.2019.

Zenchak, J. R. et al. (2018) ‘Bioluminescence-driven optogenetic activation of transplanted neural precursor cells improves motor deficits in a Parkinson’s disease mouse model’, Journal of Neuroscience Research. doi: 10.1002/jnr.24237.

Zhu, H. et al. (2016) ‘Cre-dependent DREADD (Designer Receptors Exclusively Activated by Designer Drugs) mice’, Genesis, 54(8), pp. 439–446. doi: 10.1002/dvg.22949.

